# Design and optimization of silicon-based electrokinetic microchip for sensitive detection of small extracellular vesicles

**DOI:** 10.1101/2022.08.17.504250

**Authors:** Moein Talebian Gevari, Siddharth Sourabh Sahu, Fredrik Stridfeldt, Petra Hååg, Kristina Viktorsson, Rolf Lewensohn, Alessandro Gori, Marina Cretich, Apurba Dev

## Abstract

Detection of analyte using streaming current has been previously explored using both experimental and theoretical approaches. However, little has been done to develop a viable microchip which can be exploited to deliver a sensitive, robust, and scalable biosensor device. In this study, we demonstrate the fabrication of such a device on silicon wafer using a scalable silicon microfabrication technology followed by their characterization and optimization for highly sensitive detection of small extracellular vesicles (sEVs). We show that the sensitivity of the devices, estimated using a common protein-ligand pair and sEVs significantly outperforms previous reports using the same principle. Two versions of the microchips, denoted as enclosed and open-top microchip, were developed and compared aiming to discern the importance of high-pressure measurement vs easier and better surface preparation capacity. A custom-built chip-manifold allowing easy interfacing with standard microfluidic connections was also developed. By investigating different electrical, fluidic, morphological, and fluorescence measurements, we show that while the enclosed microchip with its robust glass-silicon bonding can withstand higher pressure and thus generate higher streaming current, the open-top configuration offers several practical benefits including easy surface preparation, uniform probe conjugation, and improvement in the limit of detection (LoD). We further compare two common surface functionalization strategies and show that the proposed microchip can achieve both high sensitivity for membrane protein profiling and low LoD for sEV detection. At the optimum condition, we demonstrate that the microchip can detect sEVs reaching a LoD of 10^4^ sEV/mL, which is among the lowest in the reported microchip-based methods.

## 1. Introduction

During the past years, the principle of electrokinetic–biosensing, exploiting streaming current/potential, has been applied in the detection of a wide variety of bio-analytes including proteins and ligands [1–3], DNA [4], and extracellular vesicles (EVs) [5], thereby demonstrating a potential of the method as a generic biosensor approach. Electrokinetic biosensing relies on the electrostatic and hydrodynamic interaction at the solid-liquid interface inside a microchannel and allows for label-free detection of bio-analytes. A major benefit is its high sensitivity to the surface coverage of an analyte, which has been previously studied with inorganic particles [6] and lately, for the determination of bio-analyte concentrations [7]. Besides, the method offers several practical benefits, such as low sample consumption, simple and inexpensive device architecture, and possibility to integrate with standard microfluidic technologies for sample sorting [8], enrichment, and deliveries. These advantages have attracted further interest in the method aiming for both improved understanding and exploitation of the governing principles. We have recently demonstrated that the surface charge density and the charge contrast between the sensor surface and analyte play a major role which can be exploited to achieve a better biosensor sensitivity [9, 10]. Further, by designing an appropriate charge-labelled detection probe, we showed the possibility to develop an immunosandwich assay, thereby, extending the application of the method also to assessment of complex biofluids [9]. A multiplexed detection setup for simultaneous measurement of several bio-analytes has also been reported [11].

Clearly, the detection principle has matured in its ability to analyze bio-analytes, significantly improving in both sensitivity and specificity. An improved understanding of how the physical parameters of an analyte, such as its size and charge, influence the sensor response [12] now provides us new opportunities to design a more sensitive detector. However, the developments so far have mainly been shown using non-scalable channel design, e.g. commercial silica capillaries [10], which have limited scope to exploit many of the benefits in practical settings. An implementation of the detection principle on microfabricated channels can help to further leverage some of the key advantages of the method. This includes the design of shorter and narrower channels, scalable fabrication, improving the quality of surface oxides for a better sensitivity as well as integration of multiple channels for increasing the throughput of multiplexing. In addition, such a microfabricated sensor can open new avenues for research, e.g., new material for sensing surface, integration of fluidic actuation, exploring the benefit of nano-engineered surface etc.

In this study, we report on the fabrication and characterization of such a microchip biosensor realized by microfabrication of fluidic channels on silicon wafer. Two different designs are presented: an enclosed microchip where a glass wafer is anodically bonded with the microfabricated silicon chip, and an open-top configuration where the microchannels are created by mechanically pressing a PDMS covered glass against the silicon substrate. While both the devices show a linear and reproducible streaming current as a function of the applied pressure, the open-top chip offers several practical benefits without compromising sensitivity and LoD. Using, biotin-streptavidin pair and the EV surface tetraspanins (CD9 and CD81) expressed on small EVs (sEVs) isolated from cell culture media of a non-small cell lung cancer (NSCLC) cell line, we then demonstrate that the device can outperform the previous report on LoD using the same detection principle. We also demonstrate that surface functionalization strategies can be exploited to further improve the device performance. Thus we show that using a silane-based functionalization strategy, the microchip can achieve a LoD of 10^4^ sEV/mL when captured by Membrane Sensing Peptide (MSP) probes [13–15]. The development is expected to take the sensing principle one step closer to clinical applications.

## 2. Materials and methods

In this study, the microfluidic devices were batch processed on a silicon wafer and then diced into individual sensors chips. Each microchip consisted of four interconnected microchannels of rectangular cross-section, sharing a common inlet port. The surfaces of the microchannels were made of thermally grown silicon dioxide. The morphology and surface roughness of the devices were analyzed by scanning electron microscopy (Zeiss Leo 1530 SEM) and ZYGO optical profiler (Nexview NX2), respectively. The uniformity of the chemical functionalization was investigated by atomic force microscopy (AFM; JPK Nanowizard 3) and fluorescence analysis (Zeiss axio observer 7). The wettability of the surface was analyzed by a custom-made setup consisting of a Dino-Lite AM7115MTF camera and the images were characterized by ImageJ contact angle module. After appropriate surface cleaning and chemical functionalization, the detection sensitivity of the devices was tested with both streptavidin and sEVs isolated from cell culture media of the NSCLC H1975 cell line as previously described [16].

### 2.1. Reagents

For the study we used pure deionized water (resistivity: 18 MΩ.cm) locally produced. Phosphate buffered-saline (PBS) tablets, avidin from egg-white, streptavidin (SA) from Streptomyces avidinii, Atto-565 conjugated SA, hydrogen peroxide and ammonium hydroxide were purchased from Sigma Aldrich Sweden AB (Stockholm, Sweden). The capturing probes consisting of Poly(L-lysine)-graft-biotinylated PEG (PLL-g-PEG-biotin, referred to as PPB here after) were purchased from Nanosoft Polymers (Winston Salem, NC, USA). Silane-PEG-biotin (referred to as SPB here after) was purchased from Laysan Bio (Arab, AL, USA). Biotinylated human Anti-CD9 antibody (MEM-61; catalog no. MA1-19485) and biotinylated human anti-CD81 antibody (M38; catalog no. ab239238) were purchased from Thermofisher Scientific (Stockholm, Sweden) and Abcam (Cambridge, UK), respectively. To minimize the non-specific interaction of the sEVs with the microchip surface, pluronic (synperonic) F108 was purchased from Sigma Aldrich Sweden AB (Stockholm, Sweden). For the single EV platform, anti-CD9 antibody conjugated with VioBlue was bought from Miltenyi Biotec Norden AB (Lund, Sweden) (catalog no. 130-118-809), and anti-CD81-APC (catalog no. A87789) was purchased from Beckman Coulter, USA. All detection antibodies used on the single EV platform were monoclonal.

### 2.2. sEVs isolation, collection, and characterization

sEV isolation protocol in this study followed our previous work [16]. In short, two steps of centrifugation were performed on the cell culture media of the NSCLC cell line H1975 (ATCC® CRL-5908™, LGC Standards, Wesel, Germany). Size exclusion chromatography (SEC) on qEV original columns was performed to isolate sEVs. Finally, size and charge of the collected sEVs were characterized by nanoparticle tracking analysis (NTA, Zetaview from Particle Metrix). More details on the sEV isolation could be found in Supplementary Information (section S1).

### 2.3. Peptide design and synthesis

Synthetic Membrane Sensing Peptides (MSPs) derived from Bradykinin were used in the study for sEV capture. Unlike antibodies, MSPs show specific affinity for the highly curved lipid membrane, which can be considered a shared “epitope” for nanovesicles, making MSPs agnostic to the relative abundance of the sEV surface proteins. The peptide synthesis, e.g. Branched Peptide, followed our previous report [13]. However, modifications including a short PEG linker and a terminal Biotin handle for surface immobilization was introduced to fit our biosensor. The probe was assembled by stepwise microwave-assisted Fmoc-SPPS on a Biotage ALSTRA Initiator+ peptide synthesizer, purified by RP-HPLC and analyzed by ESI-MS as previously described [13].

### 2.4. Sensing method, measurement setup, and the microchip

The applied sensing principle is described in our previous articles [5, 11, 17]. In brief, a PBS buffer was pushed through the microchannels under hydraulic pressure to generate streaming current. A brief theoretical explanation can be found in Supplementary Information (section S2). The current was then measured using a pair of platinum (Pt) electrodes connected at both ends of the channel. The experimental setup along with the procedure is shown in Supplementary Information (section S3).

The centerpiece of the assembled system is a custom-built manifold to mount the microchip. For this purpose, an octagonal-shaped PEEK block was machined, on the center of which the microchip was placed (Figure 2a, S1). Silicone O-rings were used to ensure a leak-proof fluidic interfacing between the manifold and the microchip. Finally plastic plates were used to sandwich the microchip on the platform. An optical window was designed on the plastic holders to perform fluorescence microscopy. Finally, a custom-made PDMS twin reservoir was used on the open-top microchip to independently functionalize individual channels for the multiplexed measurements (See Supplementary Information (section S4)). The silicon and glass substrates were purchased from MicroChemicals Company (Ulm, Germany). Sylgard 184 PDMS kit was procured from Ellsworth Adhesives (Stockholm, Sweden).

**Figure 1.**
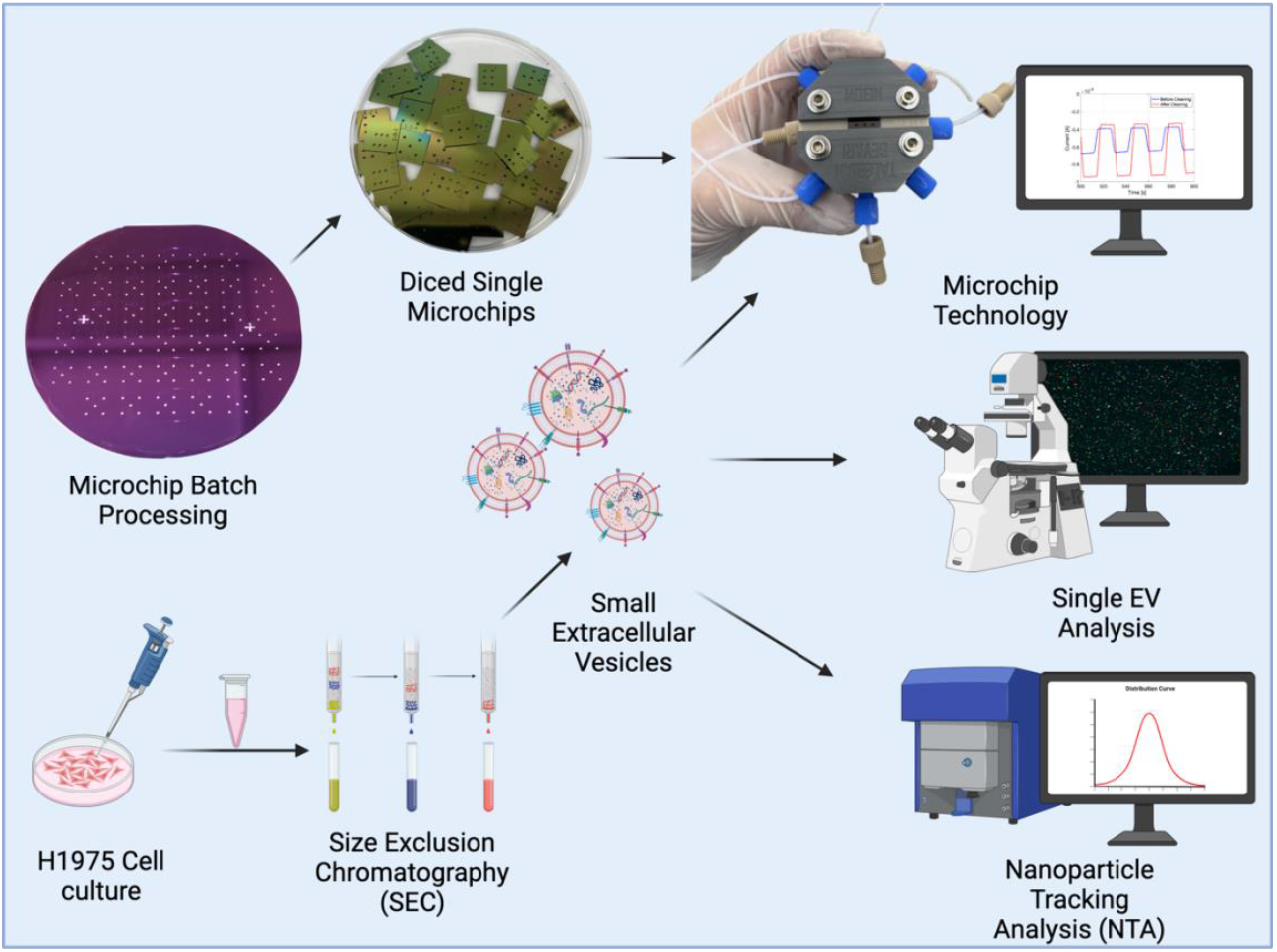
A schematic of the workflow used in this study. Microchips were batch processed on a 4-inch wafer and diced into small devices for streaming current based detection of small extracellular vesicles (sEVs) islolated from NSCLC H1975. sEVs were harvested by size exclusion chromatography (SEC). The isolated sEVs were characterized by Zetaview nanoparticle tracking analysis system for concentration, size, and zeta potential. The sEVs were further characterized using a single EV platfrom for their surface protein expression for tetraspanins CD9 and CD81.

**Figure 2.**
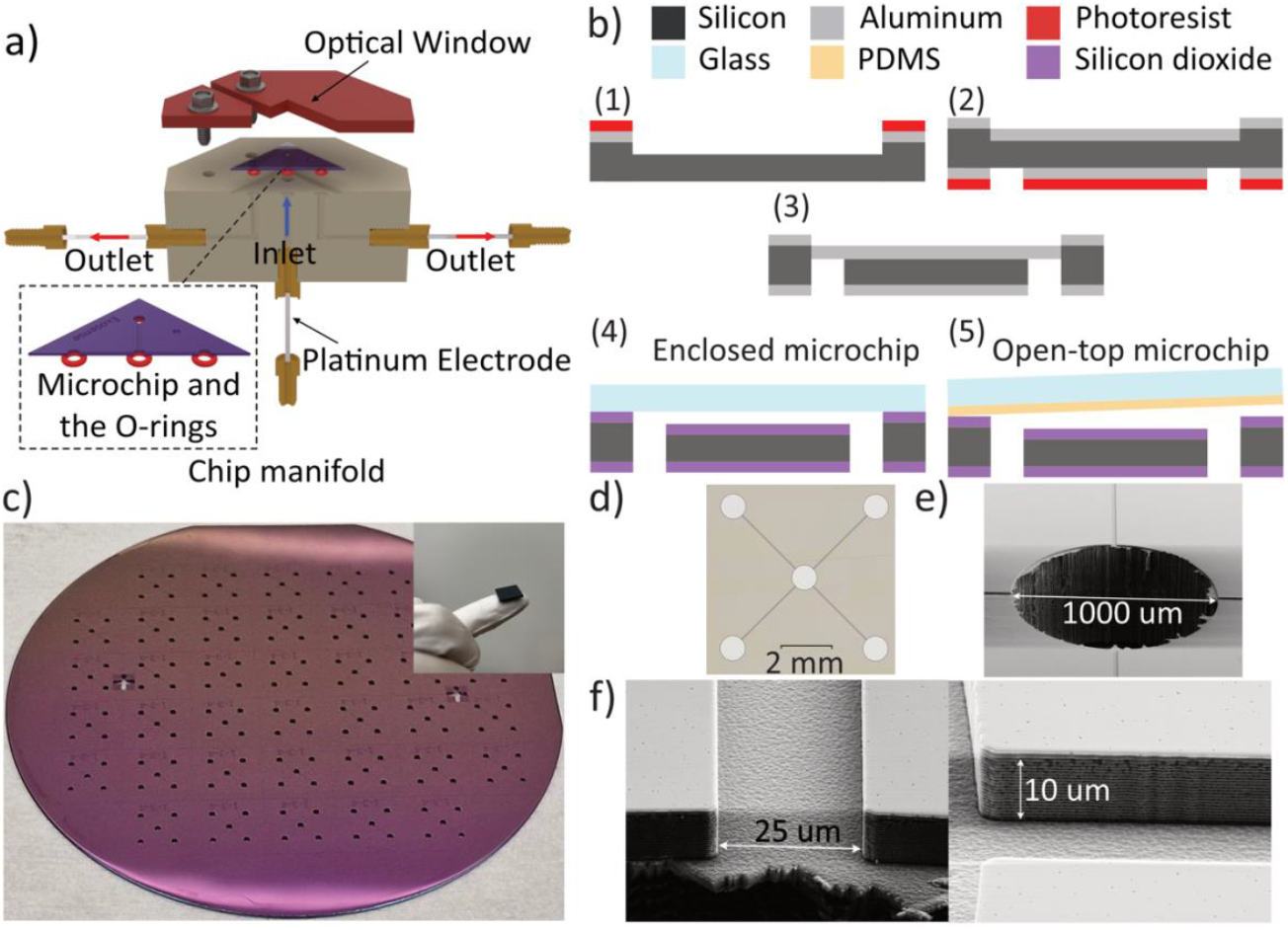
Details of the microchip fabrication and the chip manifold design. a) Schematic cross-sectional view of the chip manifold and the microchip along with the platinum electrodes and the O-rings., b) Fabrication process flow schematic 1-microchannels dry etched into silicon using a sputtered Aluminum hardmask, 2-backside optical lithography opening the fluidic ports on the Aluminum mask, 3-etch through silicon to create the inlet and outlet ports, 4-thermal oxide growth and anodic bonding with glass for enclosed chip, 5-thermal oxide growth and mechanical bonding with PDMS covered glass, c) a fully processed wafer and a single microchip on a fingertip, d) optical image of the microchip and Scanning electron microscope image of e) the inlet port and f) microchannel cross section and walls

### 2.5. Surface functionalization

Prior to the antibody/receptor immobilization, the surface of the microchannels was cleaned using an RCA1 (5:1:1 DI-Water: H_2_O_2_:NH_4_OH) solution at 88 °C for 20 minutes. Supplementary Information (section S5) demonstrates the effect of the cleaning process on the recorded streaming current. After the cleaning process the microchips were chemically functionalized either by flowing the chemicals through the channels (for enclosed chips) or simply incubating the solution (for the open-top chips). Two different surface functionalization strategies were followed in this work, *i)* a PPB-based protocol and *ii)* a SPB-based strategy. For the PPB based capture, we followed an optimized strategy as reported in our earlier work [10]. In brief, the surface of the microchannels was first coated with a thin layer of PPB by supplying an aqueous solution (0.1 mg/mL) of PPB for 15 minutes. For the biosensing, the biotinylated anti-CD9 and anti-CD81 antibodies were conjugated to the PPB coated surface using avidin as a linker molecule. The concentration of the capture antibody was 50 μg/mL in 1xPBS and was immobilized for 60 minutes. For SPB-based functionalization strategy, the surface was first treated by 1 mg/mL SPB in 95% Ethanol overnight and then washed by 95% Ethanol and DI-water. After drying, a 0.05 mg/mL solution of streptavidin in 1xPBS was used as the linker between antibodies and the surface. The antibody immobilization was identical to the step followed in the PPB case. Prior to the sensing measurements, the microchannels were treated with 0.01%w pluronic (synperonic) F108 solution for 30 minutes in order to suppress non-specific bindings.

### 2.6. Fluorescence-based single EV analysis

For comparison, fluorescence-based single EV analysis was performed following our earlier report [18]. For that, a silica coverslip was functionalized by PPB-AV using the same protocol as the microchips. Thereafter, the sEVs were incubated on the surface for 1 hour and captured electrostatically irrespective of their surface protein. The coverslip containing sEVs was then passivated for 30 minutes using 0.5 mg/mL casein to suppress non-specific binding of the fluorophore-tagged antibodies. Fluorophore tagged anti-CD9 and anti-CD81 antibodies were then incubated on the captured sEVs for 1 hour followed by washing with PBS to remove unbound antibodies. Fluorescence imaging was done with a Zeiss inverted epi-fluorescence microscope under LED excitation.

### 2.7. Microchip design and fabrication

For the fabrication, a 100 nm thick sputtered aluminum layer was used as a hard mask for etching the silicon substrate. The patterns defining the channels were lithographically generated on the surface. The aluminum mask was dry etched followed by a 10 μm deep dry etching of the silicon substrate (Figure 2b-1). Thereafter, aluminum layers were deposited on back and front sides of the substrate. A back-side photolithography followed by a dry etching of the aluminum mask (Figure 2b-2) and a deep dry etching of the silicon wafer was done (Figure 2b-3) to form the inlet and the outlet ports. Finally, a 300 nm thick layer of silicon dioxide was thermally grown on the substrate at 1000 °C.

To create the enclosed microchips, the substrate was anodically bonded to a borofloat glass wafer (Figure 2b-4). In case of the open-top microchips, a glass wafer was bonded to a 100 μm thick sheet of PDMS by plasma treatment. The PDMS covered glass was pressed against the microchips on the chip manifold to create the leak-proof fluidic path (Figure 2b-5). More details of the fabrication process flow could be found in the Supplementary Information (section S6). Figure 2c shows a fully processed wafer containing 34 chips and a single microchip (12×12 mm) on a fingertip.

The cross-sectional dimensions of the microchannels are 10 μm × 25 μm and 3 mm in length. The optical images showing four identical microchannels with a common inlet (at the center) and separate outlets as well as scanning electron microscopy images are shown in Figure 2d-f. The surface roughness of the fabricated devices was characterized using white light interferometry (WLI) method (See Supplementary Information (section S7)).

## 3. Results

Electrical and fluidic characterizations were performed to find the relative performance and optical working range of the different microchip configurations. The sensing performance of the microchips was then analyzed by comparing their detection sensitivity and LoD for targeting streptavidin and sEVs.

### 3.1. Fluidic and electrical characterization

Figure 3a shows the volumetric flowrate measured in the enclosed and open-top microchips as a function of the applied pressure. In case of the enclosed microchip, the flowrate showed a linear and highly reproducible (standard deviation, SD = ±0.7 μL/min) dependence on the upstream pressure up to 6 bar. A simple Poiseuille estimation of the flowrate is also shown in Figure 3a which demonstrates a negligible deviation of the experimental flowrate from the estimation. In comparison, the open-top microchip showed a linear dependence only up to 2 bar and then started to leak. In the latter case, the flowrate was also lower than the theoretical estimate. Thus, to ensure a leakproof measurement, the applied pressure was kept below 1.5 bar for the open-top microchips.

**Figure 3.**
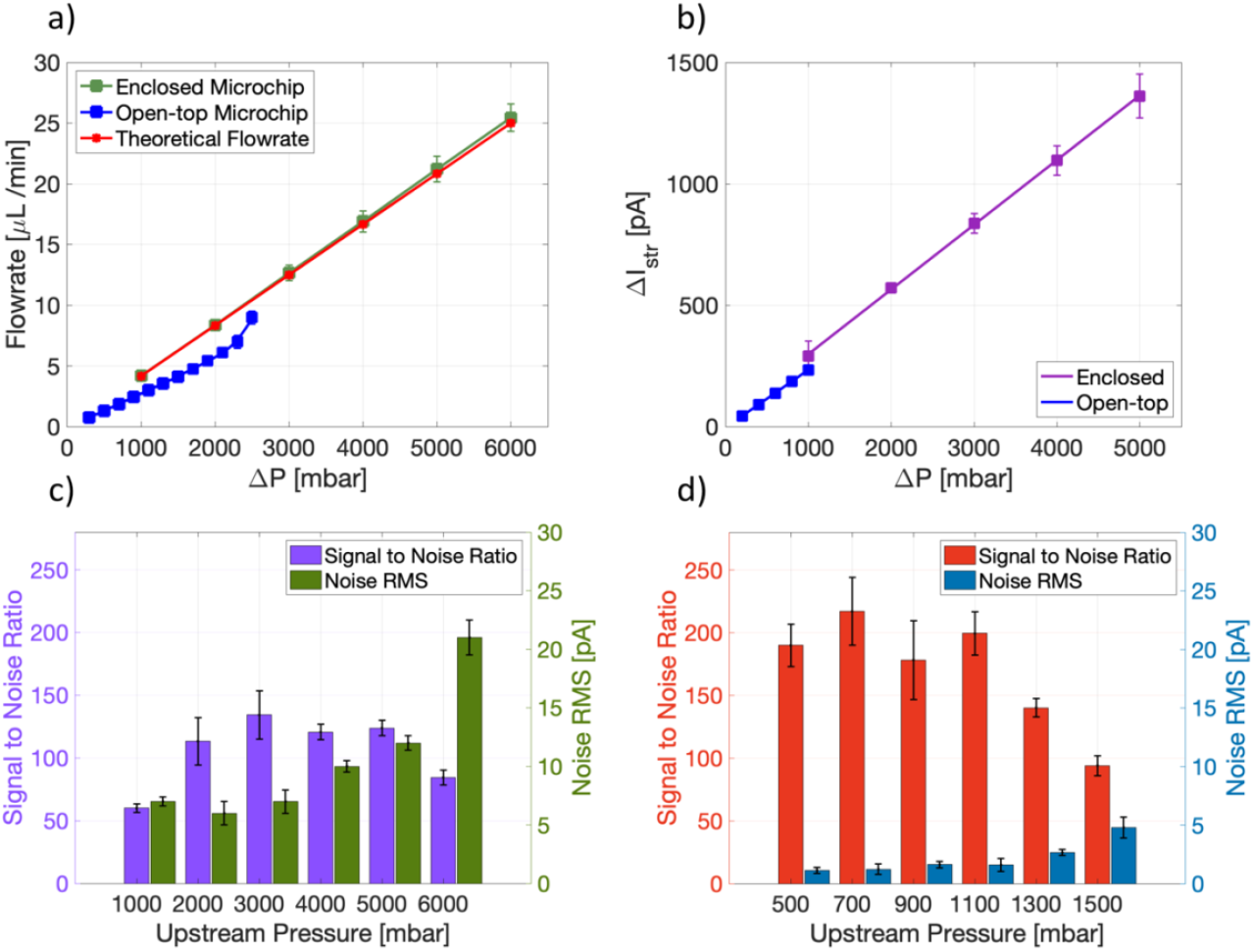
Electrical and fluidic characterization of the microchips, a) The linear relation between the flowrate and the upstream pressure for both microchips, b) The linear relation between the streaming current and the upstream pressure for both microchips, c) the Signal to Noise Ratio (SNR) and noise RMS for the enclosed microchip and d) the open-top microchip at different upstream pressures.

We then compared the streaming current magnitude and noise of the devices as a function of the applied pressure. Figure 3b shows a comparison of streaming current for the two freshly cleaned microchips. Since the geometries of the channels, are identical in both the designs, we hypothesize that the streaming current magnitude will be a measure of their relative surface qualities. It should be noted that unlike the enclosed microchip, the open-top version is confined by three active surfaces. The top surface (PDMS) does not contribute to the streaming current as much as the SiO_2_ surfaces due to the inertness of PDMS [19]. Therefore, a lower streaming current is expected. As seen, both the devices show a linear dependence of streaming current on the applied pressure, underscoring negligible effect from electrode polarization or surface conductivity [20].

Next, we compared the noise characteristics of the devices. The RMS noise was calculated at different steaming current, i.e, at different upstream pressure. The data is presented as bar plots along with the signal to noise ratio (SNR) in Figure 3c and d for the enclosed and the open-top versions, respectively and the streaming current values at constant upstream pressure are shown in Supplementary Information (section S8). We observed that the RMS noise increases with increasing streaming current. As seen, the RMS noise for the enclosed microchip remains similar (7 pA with standard deviation below ±2 pA, n=3) up to 3 bars of applied pressure but then sharply increases, reaching 21 pA at 6 bars. Similar RMS noise behavior was observed in case of the open-top microchip. The maximum RMS noise for the open-top microchip was below 5 pA, which is lower than the enclosed microchip at the same applied pressure. The comparison of SNR for the two designs clearly suggests that the open-top chip has a higher SNR at all the pressure ranges despite having a lower active surface area. Thus, for the sensing experiments, the pressure pulses were chosen to have maxima and minima at 3 bar and 1.5 bar, respectively for the enclosed microchips and 1.5 bar and 500 mbar for the open-top microchip. The noise RMS at the maxima of the pressure pulses, i.e. 7 pA and 5 pA for enclose and open microchips, respectively, were defined as the minimum detectable signal (MDS) for both devices.

### 3.2. Sensing performance of the microchips

The sensing performance of the microchips was first investigated using the biotin-streptavidin pair. To compare the performances, different concentrations of SA were detected using the PPB-based functionalization on both the microchips. Figure 4a shows the response (ΔI_s_) of the chips as a function of SA concentration. The signal from 1 nM SA was well above the MDS for both devices. The calibration plot intersecting MDS line shows a LoD of 0.57 nM and 0.45 nM for the enclosed and open-top microchip, respectively. Coefficient of determination for linear regression (R^2^) shows how well the fitted line characterized the dynamic range of the microchips in detecting SA. Clearly again, the open-top configuration shows similar performance as the enclosed version despite their differences in terms of the active surface area. Moreover, the LoD of the enclosed and the open-top microchips were lower by a factor of 3.7 and 4.9, respectively, (see Supplementary Information (section S9)) compared to the capillary-based detection used in previous studies [10, 11].

**Figure 4.**
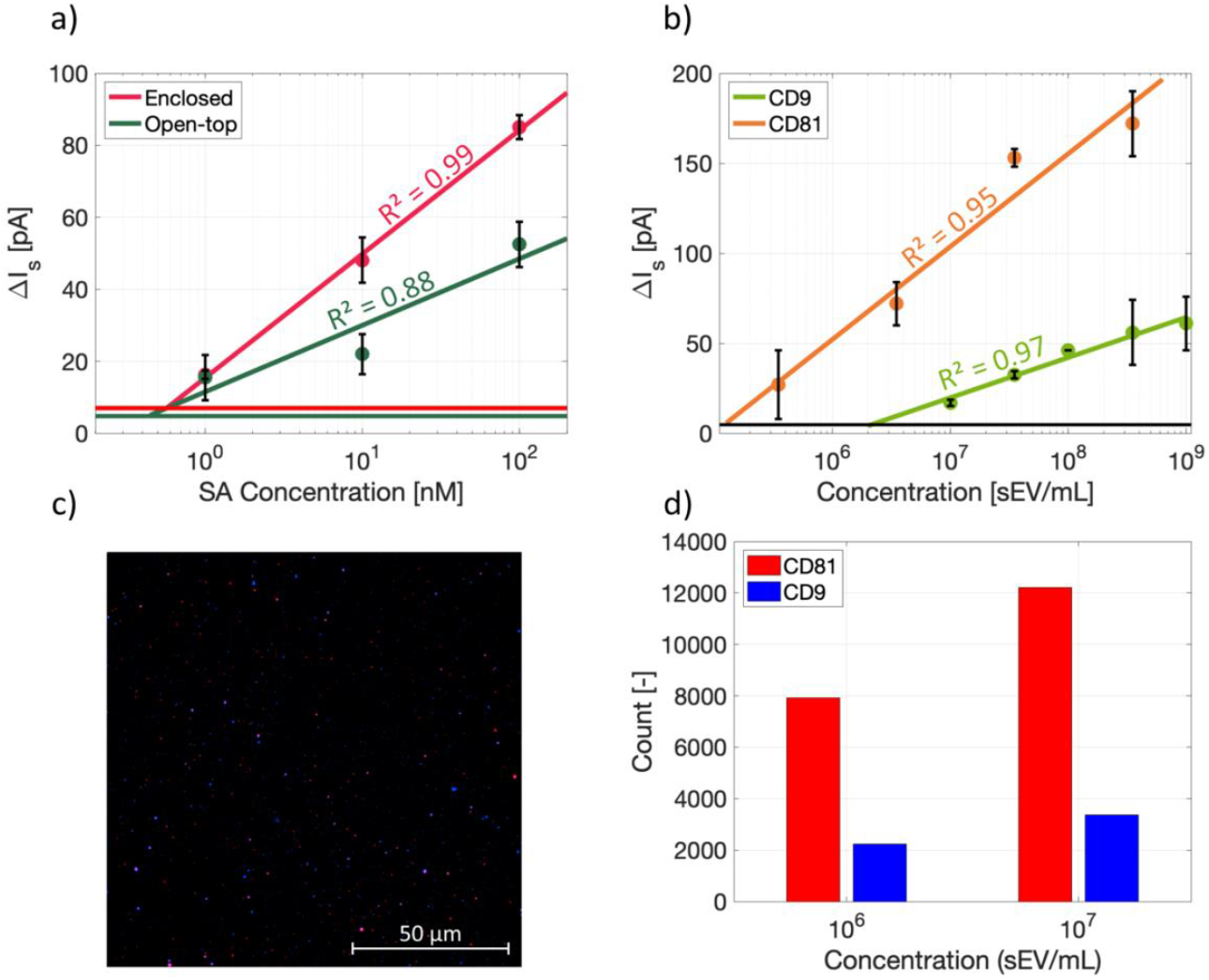
Evaluation of the sensing performances of the microchips. In a and b, the horizontal lines are corresponding minimum detectable signal and the goodness of the fit has been indicated in the plots, a) SA concentration measurements using enclosed and open-top microchips, b) Multiplexed detection of H1975 sEVs targeting CD9 and CD81 using the open-top microchip. The error bars represent SD from technical replicate measurements. c) A representative single EV fluorescence microscopy image of H1975 sEVs, with blue spots indicating sEVs expressing CD9 and red spots for CD81, d) Comparison of sEV counts expressing CD9 and CD81 for two different concentrations.

Using the transparent window design, we further performed fluorescence-based analysis of the microchip surface (Supplementary Information (section S10)) using Atto 565 conjugated SA. Furthermore, we compared the capture density and uniformity of the different designs. For this purpose, the surface of the devices was conjugated with PPB and fluorescently tagged SA, under identical conditions. The results presented in Supplementary Information (section S11) shows a non-uniform coverage of the FL-tagged SA along the microchannels in case of the enclosed microchip and a uniform coverage along the open-top device.

With clear advantage of the open-top version over the enclosed microchip, we proceeded to utilize the open-top configuration for sEV analysis. We analyzed the detection sensitivity of sEVs isolated from the cell culture media of NSCLC H1975 cells. Prior to the electrokinetic measurements, the sEVs were first characterized using a NTA for their concentration, size, and zeta potential (Supplementary Information S12). The mean diameter of the sEVs as well as their zeta potential were measured to be 153 nm and -25.83 mV, respectively. Fig. 4b shows the calibration plot depicting the signal (ΔI_s_) vs sEV concentration. The measurements were performed using the multiplexed configuration of the open-top chip targeting CD9 and CD81 transmembrane proteins for a large range of concentrations. The estimated LoD for CD9 and CD81 detection were 2.2×10^6^ sEV/mL and 1.2×10^5^ sEV/mL, respectively. The control measurements are presented in Supplementary Information (section S13). The observed LoD for CD9 is 2.2 times lower than the value reported earlier by the same sensing method on commercial capillaries [10]. Furthermore, Figure 4b suggests that the number of the captured CD81 positive sEVs are more than CD9 positive sEVs in the sample. To verify this observation, a fluorescence-based single-EV analysis was performed. A representative fluorescence image depicting single sEVs stained with VioBlue-CD9 and APC-CD81 is shown in Figure 4c. The estimated numbers of CD9- and CD81-positve sEVs in a total of 10 images as a function of concentration is shown in Figure 4d. As seen, CD81-positive sEVs were more abundant in the sample, thus, supporting the observed sensor response.

### 3.3. Enhancement of LoD

Following our previous report [10] and to further improve the LoD, we investigated if the choice of the chemical functionalization strategy can further improve the device sensitivity. We therefore compared two common chemical surface functionalization strategies involving SPB and PPB. Figure 5a shows the signal vs concentration plot of CD9 and CD81 detection for the open-top microchip using a SPB-based surface functionalization. An identical method and sample as used in Figure 4b was used here. As seen, the LoD of the open-top chip was significantly lower compared to the enclosed microchip. For better comparison, the LoD obtained with the two functionalization methods is presented in Figure 5b. As seen, SPB-based method allows to reach a LoD of 9.5 ×10^3^ sEV/mL in case of CD81 and 7.6 ×10^4^ sEV/mL for CD9, both of which are lower by a factor of 13 and 29, respectively, as compared to PPB based functionalization and a factor of 65 times lower than the capillary based method which was used for detection of CD9 before [10]. Further, the optimized surface functionalization led to very low sensor response to negative controls as shown in Supplementary Information (section S13).

**Figure 5.**
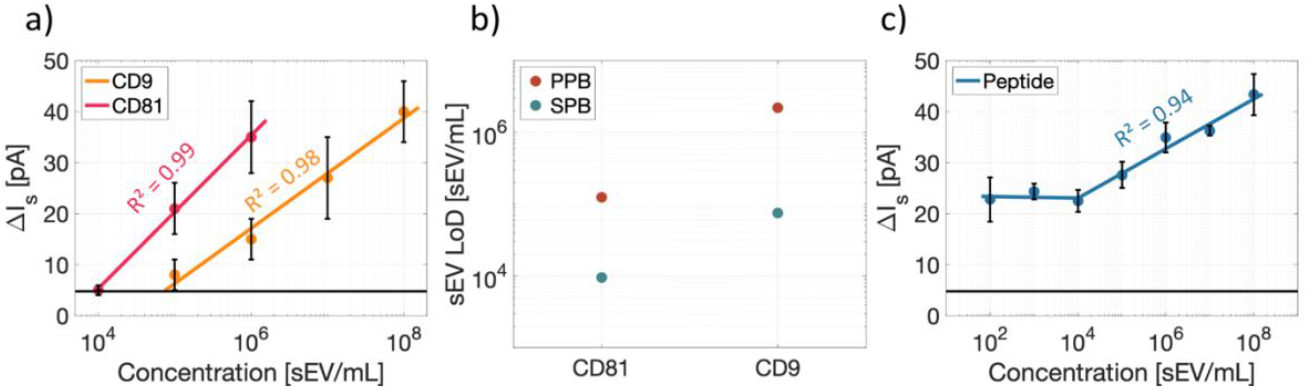
Sensing performance of the open-top microchip. *In a and c, the horizontal lines are corresponding minimum detectable signal and the goodness of the fit has been indicated in the plots*. The error bars represent SD from technical replicate measurements. a) Concentration curve for CD81 and CD9 on SPB-SA surface. b) Limit of detection comparison between PPB and SPB surfaces on the open microchip, c) Electrokinetic signal for H1975 sEVs captured by Membrane Sensing Peptides (MSPs) on SPB surface. The lowest two concentrations are out of the sensing dynamic range of the system.

Although, the analysis of sEV membrane protein expression has high clinical importance, immuno-capture may not be the best to benchmark a sensor performance. This is due to the high heterogeneity of sEVs in terms of protein expression e.g., tetraspanins [21, 22] and the relative abundance of certain sEV population. Besides, the affinity of the selected antibody may differ depending on their origin. Therefore, we further analyzed the detection sensitivity of the device using MSPs. Unlike antibody-based capture, these probes can directly bind to the sEV-membrane and thus provide a more accurate estimation of the sEV concentration. As reported earlier [13], the selected MSP can capture sEVs in the size range of 50-130 nm irrespective of their surface protein profile. Figure 5c shows the calibration curve obtained with the open-top chip and SPB based functionalization. As seen, a LoD of 10^4^ sEV/mL was obtained in that case before the sEV concentration exited the dynamic range of the sensor.

## 4. Discussion

As mentioned, the detection method has been widely studied [11, 17, 23] and improved by many research groups. In addition, the diagnostic opportunities of the method has also been explored, e.g., monitoring the efficacy of precision cancer medicine [11], albeit in a laboratory setting. Translation of these technical developments for potential use in a clinical setting with a viable microchip is the obvious next step. We present here such a microchip, which can be mass produced on silicon wafer. We used different characterization methods and bio-analysis, to addressed three main aspects: *i)* A design aspect and its influence on the device performance, *ii)* Impact of the chemical functionalization strategies on the device characteristics and *iii)* Sensitivity and the LoD of the sensors in comparison to other methods.

### Design aspect and its influence on the device performance

The advantage of silicon-based technologies for scalable fabrication is well-known [24]. Besides, SiO_2_ surface has also been widely studied for the immobilization of affinity probes [23, 25], thus justifying the selection of material and process technologies reported here. As presented in Supplementary Information (section S2), streaming current proportionally increases with the pressure difference. Hence, an obvious design choice would be a mechanically robust microchannel like the enclosed microchip which can withstand a higher pressure. As seen in Figure 3b, while ΔI_str_ expectedly increases with ΔP, it does not necessarily translate to increasing SNR (Figure 3c) for the entire range of ΔP. Since the noise RMS is roughly constant at low pressures and streaming current scales with the upstream pressure (Figure S7), the SNR increases at the beginning and then appears to reach a plateau before dropping at higher ΔP. While the mechanism behind such a noise behavior requires further study, which is beyond the scope for the present investigation, it motivated us to examine an open-top version that only works in the low ΔP region (Figure 3a and b). However, unlike the enclosed design, the open-top configuration allows full access to the sensing surface making it more convenient and compatible with different surface functionalization strategies including automated printing of affinity probes [26]. As seen in Figure S7 and 3d, the open-top configuration also produces a similar I_str_ at ΔP of 1.5 bar but higher SNR due to lower noise RMS. Given that the open-top design has 35% less active surface than the enclosed version, the result is interesting and likely indicates a higher surface activation level than the enclosed version. Besides, the LoD comparison presented in Figure 4a clearly suggests that open-top design is better suited for sensing application.

### Impact of the chemical functionalization strategies on the device characteristics

Streaming current-based approach also critically depends on the choice of linker molecules that binds affinity probes to the surface. This is primarily due the influence of surface roughness and charge contrast between an analyte and the surface, as reported earlier [10]. To further improve the performance of the device, we investigated the relative influence of PPB vs SPB coated surface. SPB based strategy allowed us to lower the LoD by more than a factor of 10 over the PPB coated surface (Figure 5b). To investigate this further, we performed AFM and contact angle measurement as presented in Figure 6. For this, we used two silica coverslips which were functionalized by PPB and SPB. The mean roughness for the SPB and PPB coated surface was 0.8 nm and 1.4 nm, respectively (Figure 6a). It is known that the surface roughness in the order of the Debye length can reduce the influence of particle adsorption on the generated streaming current [27]. This may explain the higher LoD for PPB coated surface. Furthermore, the contact angle measurement shows a 14-degree difference between PPB and SPB surfaces indicating that the PPB surface was more hydrophobic (Figure 6b). This means that the ions in the electric double layer will likely slip faster on the surface and generate more absolute value of the streaming current [28, 29]. Slide angle measurements comparing the friction force between the liquid and the surface were also carried out, Supplementary Information (section S14), and supported this claim. Higher streaming current generation on the PPB surface would lead to a steeper slope in the concentration curves by increasing the sEV concentration. This is evident when comparing the same markers of the sEVs on both PPB and SPB surfaces.

**Figure 6.**
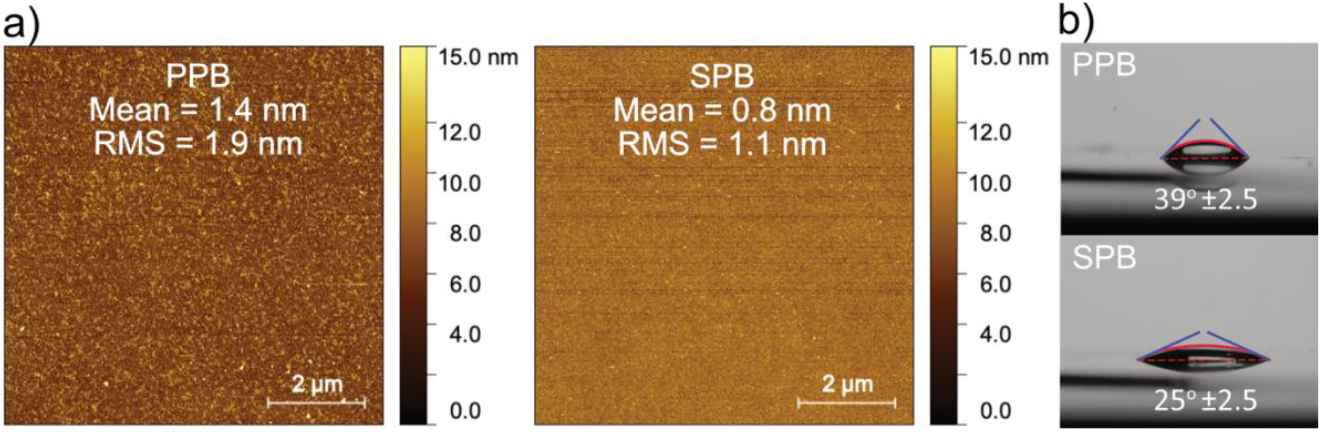
Comparison of the surface roughness and hydrophobicity of the PPB and SPB surfaces - a) AFM images of SPB and PPB surfaces comparing the mean and RMS of the roughness. The higher roughness of the PPB surface could be a possible reason for its failure to detect very low concentrations of analytes. b) Contact angle comparison between PPB and SPB surfaces, demonstrating that the PPB surface is more hydrophobic.

### Sensitivity and the LoD of the sensors in comparison to other methods

The data presented in Figure 4 and 5 clearly suggest that the fabricated microchip offers a far better LoD compared to the previous reports using the same principle [10]. A key advantage of the method stems from its dependence on the size of the analyte, an aspect which we have previously investigated using both theoretical and experimental approaches [12]. Thus, analytes like sEVs are suitable candidates for the method allowing to achieve an extremely low detection limit. A major challenge is, however, the heterogeneity of sEV samples with respect to their membrane protein composition. Therefore, to ensure similar conditions, the comparison has been done with our previous studies only where identical sample and antibody have been used. This is however difficult to maintain while comparing among different methods reported by different groups. In this context, MSPs may be suitable alternative as they are able to enrich small vesicles on the basis of specific membrane biophysical traits, opposed to the pre-selection of sEV sub-populations introduced by the use of antibodies [13, 15]. As presented in Figure 5c, a LoD of 10^4^ sEV/mL could be achieved using such peptides. Although a direct comparison of LoD among different devices is difficult, a qualitative assessment may still be possible. Table 1 shows the state-of-the-art techniques and their reported LoD compared with the present work. Clearly, the proposed method is among the best performing methods.

**Table 1.** Limit of detection and linear range reported in the literature targeting EVs on different platforms.

| Platform | LoD [EV/mL] | Linear range [decades] | Reference |
| --- | --- | --- | --- |
| This work | $1 \times 10^4$ | 4 | N/A |
| iMEX | $3 \times 10^4$ | 4 | [30] |
| Covalent organic framework | $1.6 \times 10^5$ | 5 | [31] |
| Colorimetry | $2.76 \times 10^6$ | 1 | [32] |
| Colorimetry | $5.2 \times 10^8$ | 1 | [33] |
| Electrochemiluminescence | $7.41 \times 10^4$ | 3 | [34] |
| Electrochemistry | $2.09 \times 10^4$ | 7 | [35] |
| Electrochemistry | $2 \times 10^5$ | 4 | [36] |

### Conclusion

In conclusion, we have demonstrated the fabrication and characterization of a novel microchip based electrokinetic biosensor. The devices were fabricated on a silicon platform using a scalable process technology. We investigated different aspects of the microchip development including the design considerations, electrical and fluidic behavior and their relative performance to biosensing, thus giving a practical and necessary guideline to develop and implement such a biosensor. A custom-built chip manifold was also designed for an easy interfacing of standard fluidic connectors with the microchips and for a leak-free flow of electrolyte. The sensitivity and LoD of the microchips were compared with previous reports demonstrating their superior performance. Particularly, for the detection of sEVs, we demonstrated that the proposed microchip could offer over 60x lower LoD than the previous reports using the same principle.

## Acknowledgement

This study was supported by grants from the Swedish Research Council (grant no. 2016-05051 and 2018-06228), the Erling Persson Family Foundation, Stockholm Cancer Society (contract no. 201202, 191293, 221212 and 221383), the Swedish Cancer Society (contract no. CAN2021/1469) and Stockholm County Council (contract no. 909121 and 750032). We acknowledge Myfab Uppsala for providing facilities and experimental support. Myfab is funded by the Swedish Research Council (2019-00207) as a national research infrastructure. Work was partially funded by the European Union through Horizon 2020 research and innovation program under grant agreement No. 951768 (project MARVEL).

## References

[1] C. Glad, K. Sjödin, and B. Mattiasson, “Streaming potential—a general affinity sensor,” Biosensors, vol. 2, no. 2, pp. 89–100, 1986.

[2] A. Dev et al., “Electrokinetic effect for molecular recognition: A label-free approach for real-time biosensing,” Biosens. Bioelectron., vol. 82, pp. 55–63, 2016.

[3] J. O. Lee, N. Choi, J.-W. Lee, S. Song, and Y.-P. Kim, “Rapid electrokinetic detection of low-molecular-weight thiols by redox regulatory protein-DNA interaction in microfluidics,” Sensors Actuators B Chem., vol. 336, p. 129735, 2021.

[4] Y. Li, S. N. Lai, and B. Zheng, “A microfluidic streaming potential analyzer for label-free DNA detection,” Sensors Actuators B Chem., vol. 259, pp. 871–877, 2018.

[5] S. Cavallaro et al., “Label-free surface protein profiling of extracellular vesicles by an electrokinetic sensor,” ACS sensors, vol. 4, no. 5, pp. 1399–1408, 2019.

[6] D. Lupa, M. Oćwieja, N. Piergies, A. Baliś, C. Paluszkiewicz, and Z. Adamczyk, “Gold nanoparticles deposited on silica microparticles-Electrokinetic characteristics and application in SERS,” Colloid Interface Sci. Commun., vol. 33, p. 100219, 2019.

[7] H. Qi et al., “Rapid detection of trace Cu2+ using an l-cysteine based interdigitated electrode sensor integrated with AC electrokinetic enrichment,” Electrophoresis, vol. 40, no. 20, pp. 2699–2705, 2019.

[8] S. D. Ibsen et al., “Rapid isolation and detection of exosomes and associated biomarkers from plasma,” ACS Nano, vol. 11, no. 7, pp. 6641–6651, 2017.

[9] S. S. Sahu et al., “Electrokinetic sandwich assay and DNA mediated charge amplification for enhanced sensitivity and specificity,” Biosens. Bioelectron., vol. 176, p. 112917, 2021.

[10] S. S. Sahu et al., “Exploiting Electrostatic Interaction for Highly Sensitive Detection of Tumor-Derived Extracellular Vesicles by an Electrokinetic Sensor,” ACS Appl. Mater. Interfaces, 2021.

[11] S. Cavallaro et al., “Multiplexed electrokinetic sensor for detection and therapy monitoring of extracellular vesicles from liquid biopsies of non-small-cell lung cancer patients,” Biosens. Bioelectron., vol. 193, 2021, doi: 10.1016/j.bios.2021.113568.

[12] S. S. Sahu, C. Stiller, S. Cavallaro, A. E. Karlström, J. Linnros, and A. Dev, “Influence of molecular size and zeta potential in electrokinetic biosensing,” Biosens. Bioelectron., vol. 152, p. 112005, 2020.

[13] A. Gori et al., “Membrane-binding peptides for extracellular vesicles on-chip analysis,” J. Extracell. Vesicles, vol. 9, no. 1, p. 1751428, 2020.

[14] B. Benayas et al., “Proof of concept of using a membrane-sensing peptide for sEVs affinity-based isolation,” 2023.

[15] A. Gori et al., “Addressing heterogeneity in direct analysis of Extracellular Vesicles and analogues using Membrane-Sensing Peptides as Pan-Affinity Probes,” bioRxiv, pp. 2012–2023, 2023.

[16] C. Stiller et al., “Detection of tumor-associated membrane receptors on extracellular vesicles from non-small cell lung cancer patients via immuno-PCR,” Cancers (Basel)., vol. 13, no. 4, p. 922, 2021.

[17] S. S. Sahu et al., “Multi-marker profiling of extracellular vesicles using streaming current and sequential electrostatic labeling,” Biosens. Bioelectron., vol. 227, p. 115142, 2023.

[18] S. Cavallaro et al., “Multiparametric profiling of single nanoscale extracellular vesicles by combined atomic force and fluorescence microscopy: correlation and heterogeneity in their molecular and biophysical features,” Small, vol. 17, no. 14, p. 2008155, 2021.

[19] B. K. Gale et al., “Low-cost MEMS technologies,” 2016.

[20] D. C. Martins, V. Chu, D. M. F. Prazeres, and J. P. Conde, “Streaming currents in microfluidics with integrated polarizable electrodes,” Microfluid. Nanofluidics, vol. 15, no. 3, pp. 361–376, 2013.

[21] R. R. Mizenko et al., “Tetraspanins are unevenly distributed across single extracellular vesicles and bias sensitivity to multiplexed cancer biomarkers,” J. Nanobiotechnology, vol. 19, no. 1, p. 250, 2021.

[22] C. Han et al., “Single-vesicle imaging and co-localization analysis for tetraspanin profiling of individual extracellular vesicles,” J. Extracell. Vesicles, vol. 10, no. 3, p. e12047, 2021.

[23] D. C. Martins, V. Chu, D. M. F. Prazeres, and J. P. Conde, “Electrical detection of DNA immobilization and hybridization by streaming current measurements in microchannels,” Appl. Phys. Lett., vol. 99, no. 18, p. 183702, 2011.

[24] J. D. Plummer and P. B. Griffin, Integrated Circuit Fabrication: Science and Technology. Cambridge University Press, 2023.

[25] A. J. de Jesus and H. Yin, “Computational design of membrane curvature-sensing peptides,” Comput. Protein Des., pp. 417–437, 2017.

[26] A. J. Summers et al., “Optimization of an antibody microarray printing process using a designed experiment,” ACS omega, vol. 7, no. 36, pp. 32262–32271, 2022.

[27] M. L. Ekiel-Jezewska, Z. Adamczyk, and J. Blawzdziewicz, “Streaming current and effective ζ-potential for particle-covered surfaces with random particle distributions,” J. Phys. Chem. C, vol. 123, no. 6, pp. 3517–3531, 2019.

[28] P. Karan, J. Chakraborty, and S. Chakraborty, “Electrokinetics over hydrophobic surfaces,” Electrophoresis, vol. 40, no. 5, pp. 616–624, 2019.

[29] E. Donath and A. Voigt, “Streaming current and streaming potential on structured surfaces,” J. Colloid Interface Sci., vol. 109, no. 1, pp. 122–139, 1986.

[30] S. Jeong, J. Park, D. Pathania, C. M. Castro, R. Weissleder, and H. Lee, “Integrated magneto– electrochemical sensor for exosome analysis,” ACS Nano, vol. 10, no. 2, pp. 1802–1809, 2016.

[31] M. Wang et al., “Detection of colorectal cancer-derived exosomes based on covalent organic frameworks,” Biosens. Bioelectron., vol. 169, p. 112638, 2020.

[32] R. Vaidyanathan et al., “Detecting exosomes specifically: a multiplexed device based on alternating current electrohydrodynamic induced nanoshearing,” Anal. Chem., vol. 86, no. 22, pp. 11125–11132, 2014.

[33] Y. Xia et al., “A visible and colorimetric aptasensor based on DNA-capped single-walled carbon nanotubes for detection of exosomes,” Biosens. Bioelectron., vol. 92, pp. 8–15, 2017.

[34] B. Qiao et al., “An electrochemiluminescent aptasensor for amplified detection of exosomes from breast tumor cells (MCF-7 cells) based on G-quadruplex/hemin DNAzymes,” Analyst, vol. 144, no. 11, pp. 3668–3675, 2019.

[35] S. Wang et al., “Aptasensor with expanded nucleotide using DNA nanotetrahedra for electrochemical detection of cancerous exosomes,” ACS Nano, vol. 11, no. 4, pp. 3943–3949, 2017.

[36] X. Doldán, P. Fagúndez, A. Cayota, J. Laíz, and J. P. Tosar, “Electrochemical sandwich immunosensor for determination of exosomes based on surface marker-mediated signal amplification,” Anal. Chem., vol. 88, no. 21, pp. 10466–10473, 2016.

