## Supplementary Information for "Design and optimization of silicon-based electrokinetic microchip for sensitive detection of small extracellular vesicles"

### S1. sEVs isolation, collection, and characterization

The sEVs used in this study was harvested from cell culture media of the non-small cell lung cancer (NSCLC) cell line H1975 (ATCC® CRL-5908™, LGC Standards, Wesel, Germany) essentially as previously been described [1]. Thus, EVs were coming from RPMI 1640 medium supplemented with fetal bovine serum (FBS) that were depleted of endogenous exosomes. The sEVs were isolated using size-exclusion chromatography (SEC) on qEV original columns (Izon Science, Oxford, UK) in the same manner as described in Stiller et al., [1]. The particle size and zeta potential were characterized by nanoparticle tracking analysis (NTA, Zetaview from particle Metrix). For these analyses the sEVs samples were diluted 1:100 in filtered PBS. The sample was injected in 100 µL portions for three times and the cell temperature was maintained at 24° C. The size analysis was done on eleven positions in the sample cell while for the zeta potential a two-position measurement was performed. The CD9 expression on the used sEVs were not checked by western blot specifically but is expressed in sEVs when the SEC isolation has been used as previously been reported [2].

### S2. Theoretical background

The potential difference of the ions and electrons on the solid/electrolyte interface creates the electric double layer (EDL). A pressure gradient between the two ends of the microchannel pushes the electrolyte along the charged surface inducing streaming current ($I_s$) that can be defined as the charge flowing through the interface (S) per unit of time (Eq. (1)) [4].

$$I_s = \iint_S \rho_e V . dS \tag{1}$$

where $\rho_e$ and V are electric charge density and fluid velocity, respectively. Governing the charge density by Poisson-Boltzmann equation and the laminar flow in the microchannels by Stokes equations, one could express streaming current as in eq. (2) [4]:

$$I_s = \varepsilon\varepsilon_0 \frac{A}{\eta} \frac{P}{L} \zeta \tag{2}$$

where $\epsilon\epsilon_0$ and η refer to the permittivity of the electrolyte and dynamic viscosity, respectively, and L, P, and A refer to the length, upstream pressure, and cross-sectional area of the microchannel, respectively. Finally, ζ corresponds to the interface potential known as zeta potential which is the property of the surface.

As in eq. (1) and (2), the streaming current flowing inside a microchannel depends on the upstream pressure and the surface area contributing to the streaming current generation. In addition, a uniform and dense charge distribution on the interface could increase ζ and affect the induced streaming current.

### S3. Experimental setup

A schematic of the experimental setup consisted of a high-pressure pure nitrogen gas capsule regulated by an Elveflow OB1 flow controller coupled with a thermal time of flight (TOF) flow sensor (Elveflow, MSF3) to measure the flowrate during the experiments. The electrolyte was hydraulically pushed by the nitrogen gas into the system through PEEK tubing and microfluidic connections that were procured from Darwin Microfluidics (Paris, France). A continuous train of pressure pulses with a duration of 30 s, was used to perform a two-point streaming current measurement. The resulting streaming current pulses ($\Delta I_{str}$) were measured by a Keithley picoammeter (model no. 2636A) as a function of time, thus, constituting a baseline measurement while the pressure pulses (ΔP) were applied directly by the pressure regulator. The measurements in this study involved recording the initial streaming current baseline after the immobilization of the capturing probes ($\Delta I_{str1}$) and the final baseline after the injection of the target ($\Delta I_{str2}$). The signal reported in this work is the difference between the two baselines denoted as $\Delta I_s$ (=$\Delta I_{str2}$ - $\Delta I_{str1}$). The incubation of sEVs was done in 1x PBS to resemble physiological conditions, whereas both the baselines were measured in 0.1x PBS to reduce the charge screening. Custom made Labview and MATLAB scripts were used to record and analyze the data, respectively. The noise-sensitive components of the experimental setup were placed in a Faraday cage. Suitable holes were drilled in the chip manifold with the same diameter as the inlet/outlet ports on the microchips. To eliminate the leakage from the system, silicone rubber O-rings, purchased from Apple Rubber Inc. (Lancaster, NY, USA), with 1 mm inner diameter were placed between the microchip and the manifold.

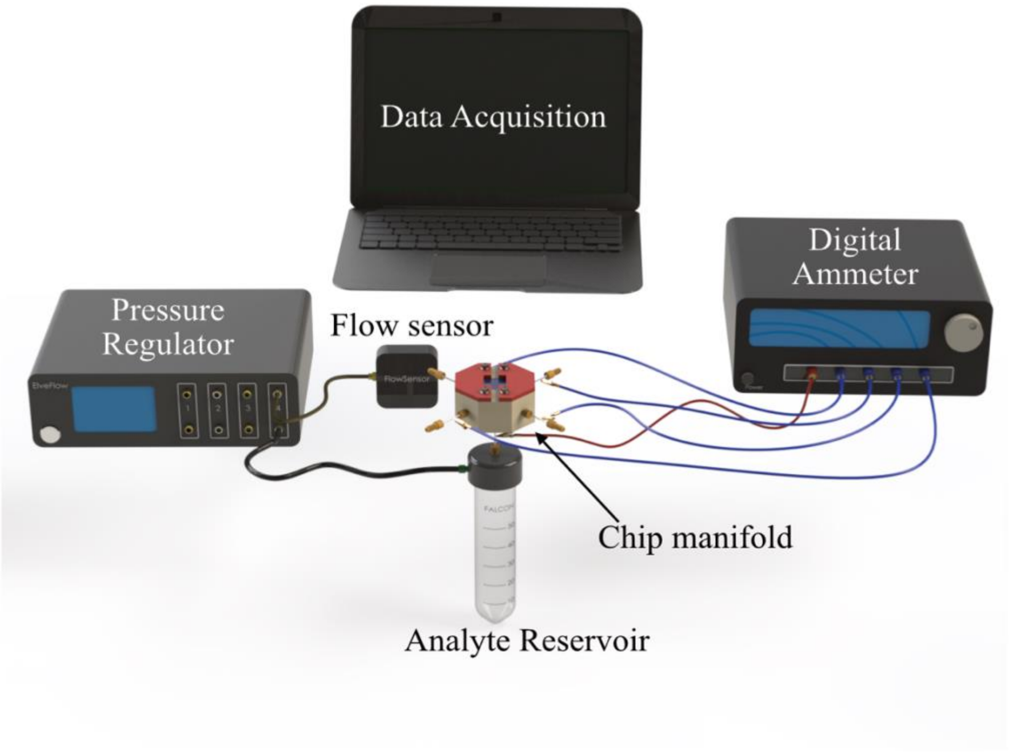


Figure *S1 – The schematic of the experimental setup consisting of the chip manifold, Platinum electrodes, digital ammeter, pressure regulator, flow sensor, analyte reservoir, and data acquisition system*

**S4. Twin reservoir and agitating platform**

A PDMS twin reservoir was casted on an in-house designed PEEK mold. Then, it was inserted on the open-top microchip to isolate two microchannels from the other two and facilitate the multiplexed measurements. Figure S2 shows the schematic of the mold and the PDMS reservoir along with the agitating platform for multiplexed antibody incubation on the microchip.

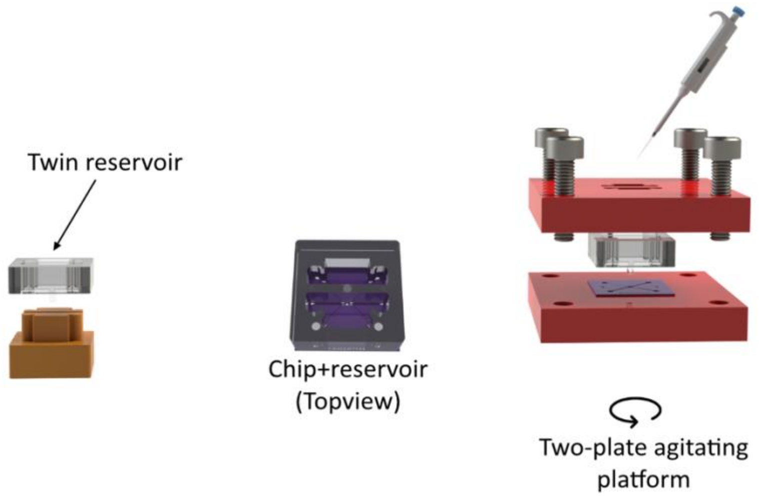


Figure *S2 – The PDMS twin reservoir molding and insertion on the chip along with the agitating platform*

To demonstrate the performance of the twin reservoir, a microchip was functionalized by PPB following the protocol explained in the manuscript. Thereafter, one side of microchip was functionalized by FL-tagged SA using the PDMS twin reservoir and the agitating platform. The FL-microscope images in Figure S3 shows the clear distinction in the signal on two sides. A border between two sides of the microchip (top) and two different microchannels (bottom) are shown in the figure.

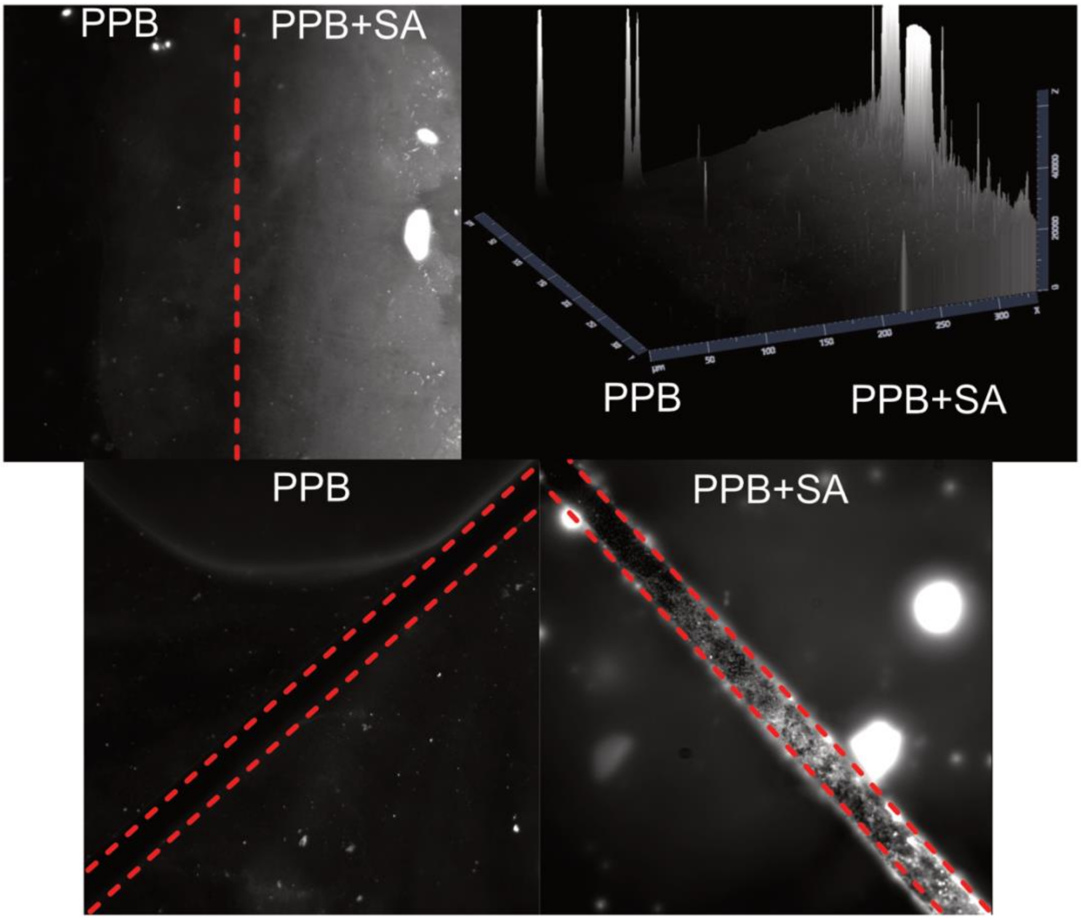


*Figure S3 – the performance of the twin reservoir. The border between the SA covered and the clear area (top), a PPB covered microchannel beside a SA covered microchannel (bottom)*

### S5. Cleaning effect

A bare microchip after the microfabrication was used to measure the baseline. Figure S4 shows the $\Delta I_{str}$ for a pressure pulse between 1.5 bar and 3 bar. Then the microchip was cleaned following the protocol explained in the manuscript and the same measurement was repeated. The effect of the cleaning on the recorded baseline is shown in Figure S4.

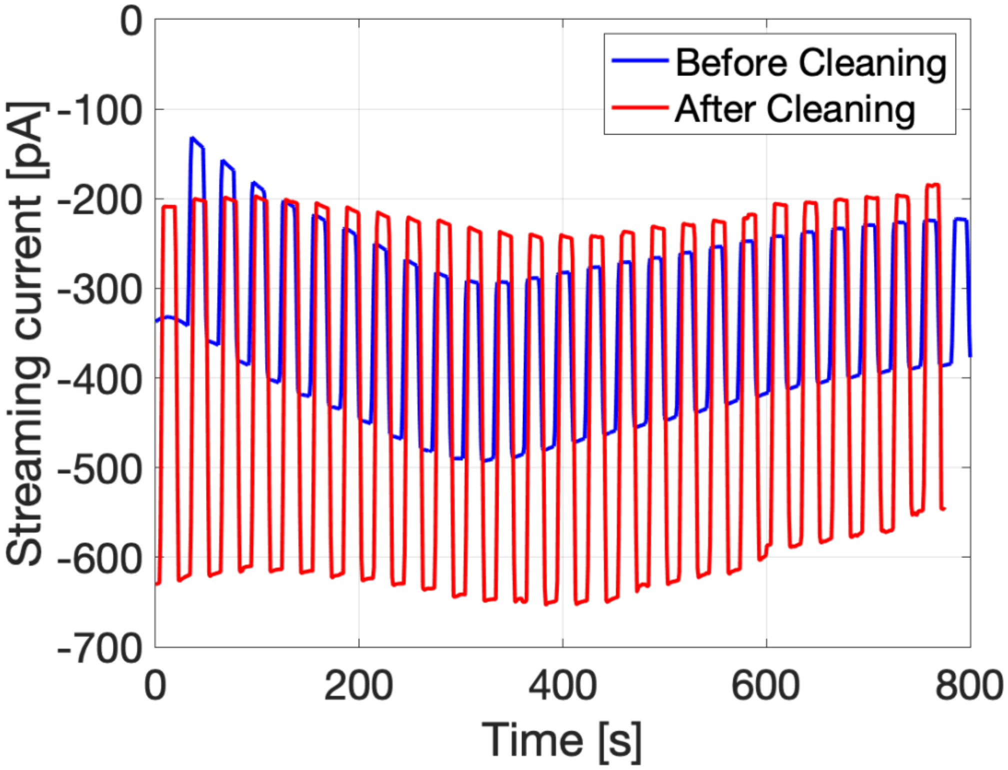


Figure *S4 – Recorded streaming current before and after cleaning*

#### S6. Fabrication process flow

A detailed fabrication process flow is shown in Figure S5.

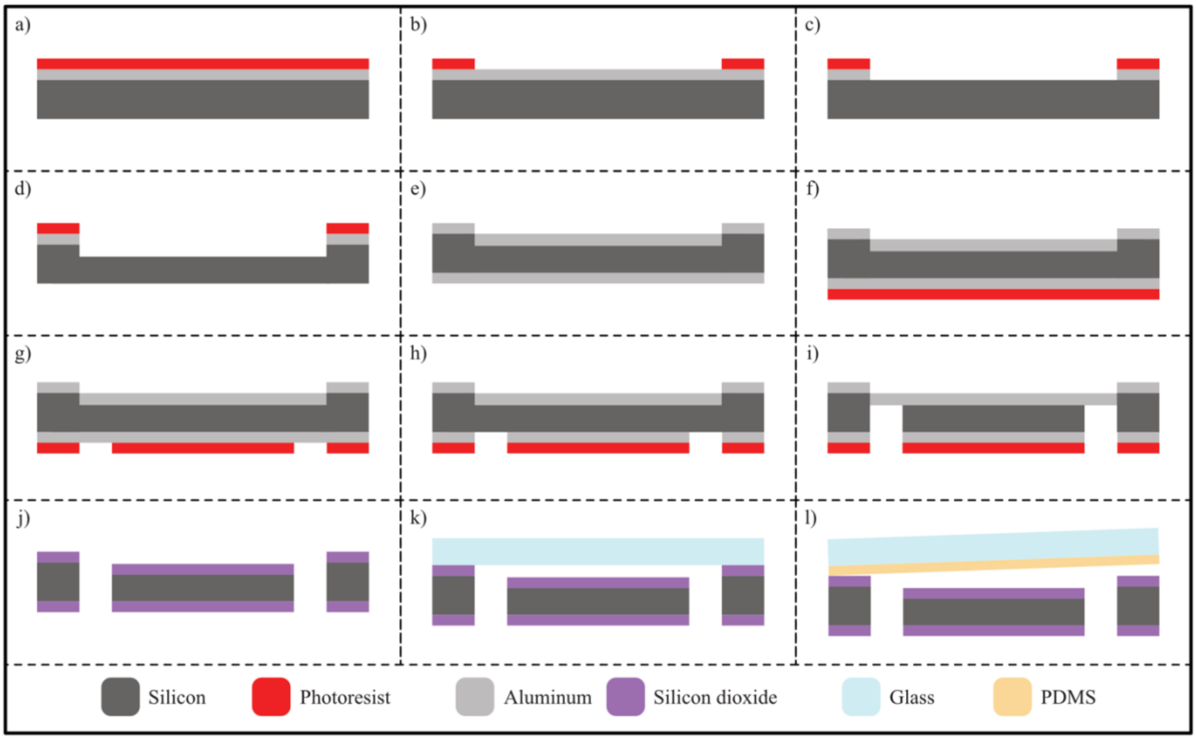


*Figure S5 – Cross sectional view of the fabrication process flow, a) silicon substrate covered by aluminum and photoresist, b) the developed photoresists after the first lithography, c) dry etched aluminum layer and open silicon surface, d) dry etched silicon forming the structure of the microchip, e) redeposition of aluminum on both front and back sides, f) spun coated photoresist for the second lithography on the back side, g) developed photoresist after the second lithography, h) dry etched aluminum mask, i) dry etched silicon forming the inlet and outlet ports, j) thermally grown silicon dioxide acting as the active surface of the sensor, k) anodically bonded glass ca, l) PDMS covered glass on the open microchip*

### S7. White Light Interferometry characterization of the microchip surface

To characterize the surface roughness of the microchips using white light interferometry, an open microchip was cleaned using RCA 1 cleaning solution and the structure was characterized by a ZYGO optical profiler as shown in Figure S6. The inset of this figure shows representative individual roughness elements analyzed on the surface. A total number of about 340K elements were analyzed and the centerline average was 9 nm.

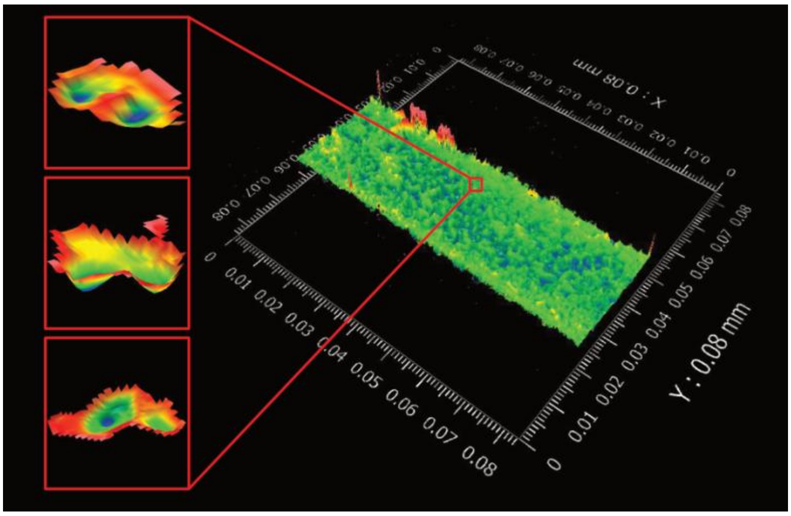


Figure *S6 – White Light Interferometry image of the microchip and the individual roughness elements*

### S8. Streaming current values at constant upstream pressure

To find the noise RMS, the upstream pressure was kept constant and streaming current was measured. Figure S7 shows the streaming current values.

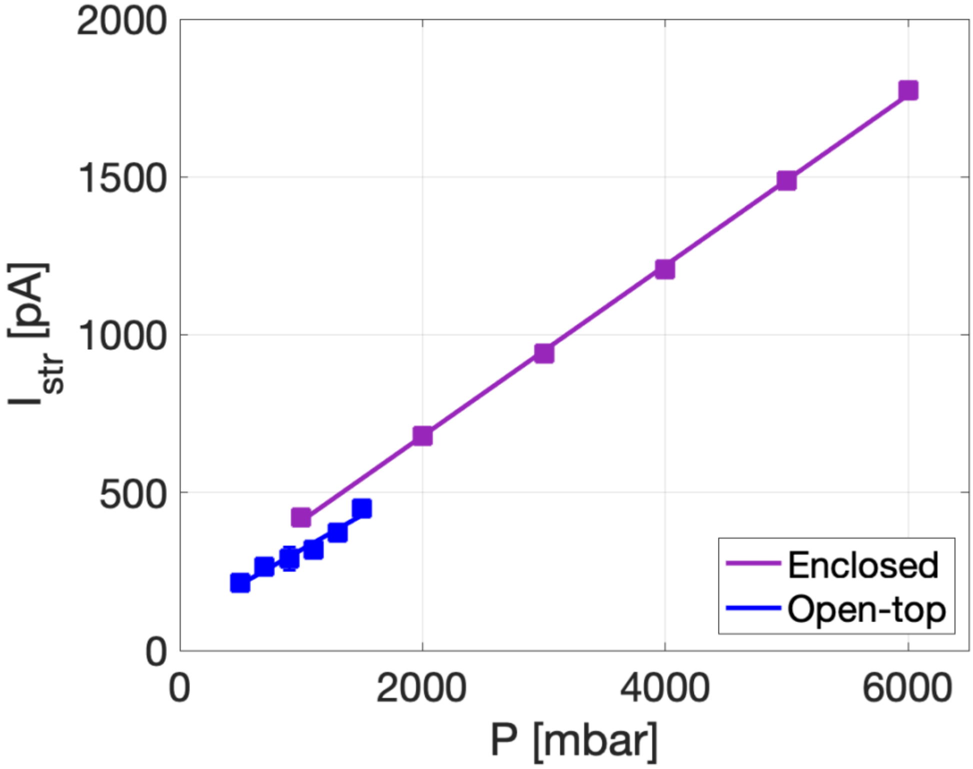


Figure *S7 – streaming current baseline of enclosed and open-top microchips at different upstream pressures.*

### S9. Streptavidin detection using different devices

The performances of both enclosed and open-top microchips were compared with commercial silica capillary tubes. The concentration curve in Figure S8 shows a significant LoD enhancement for both the microchips as compared to the silica capillaries.

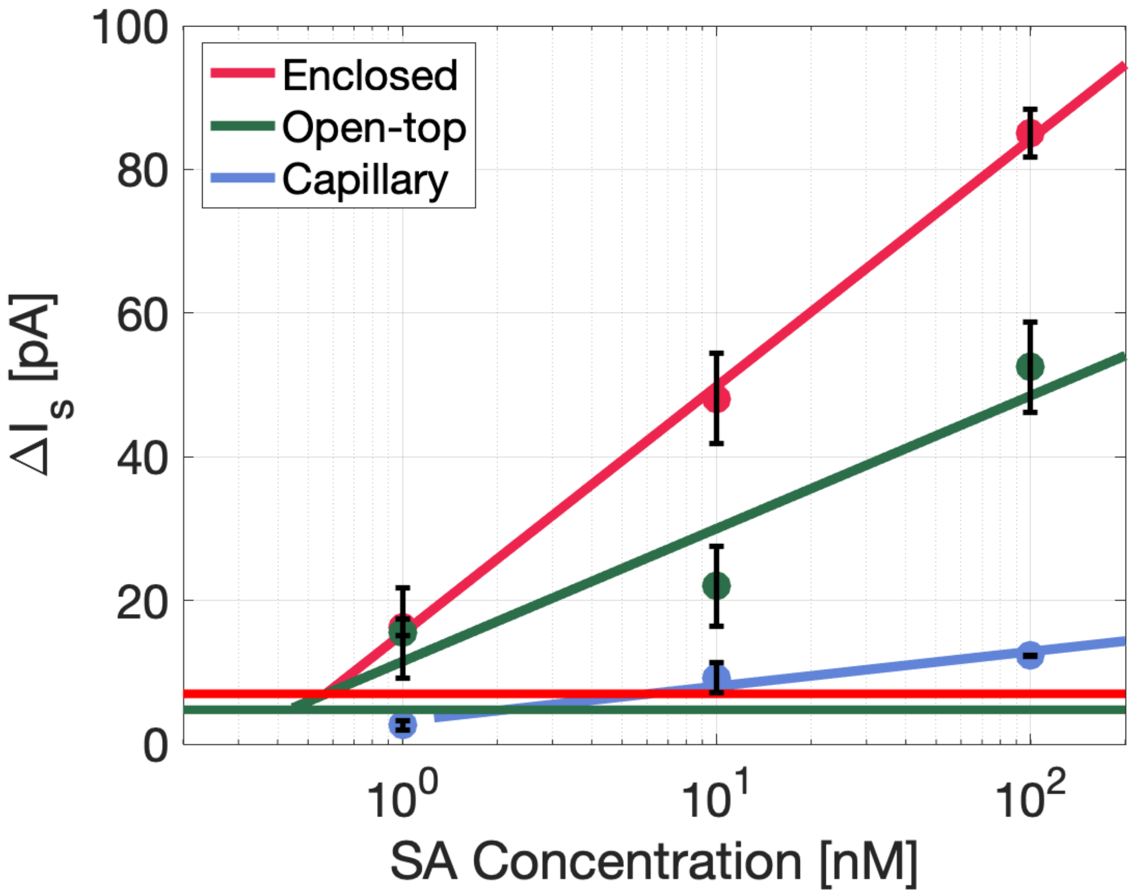


Figure *S8 – Concentration curve for SA detection on PPB surface using enclose microchip, open-top microchip, and commercial capillary tubes*

**S10. SA FL enclosed**

The simultaneous electrokinetic and fluorescent measurements on the enclosed microchip through the optical window in detecting FL-tagged SA is shown in Figure S9

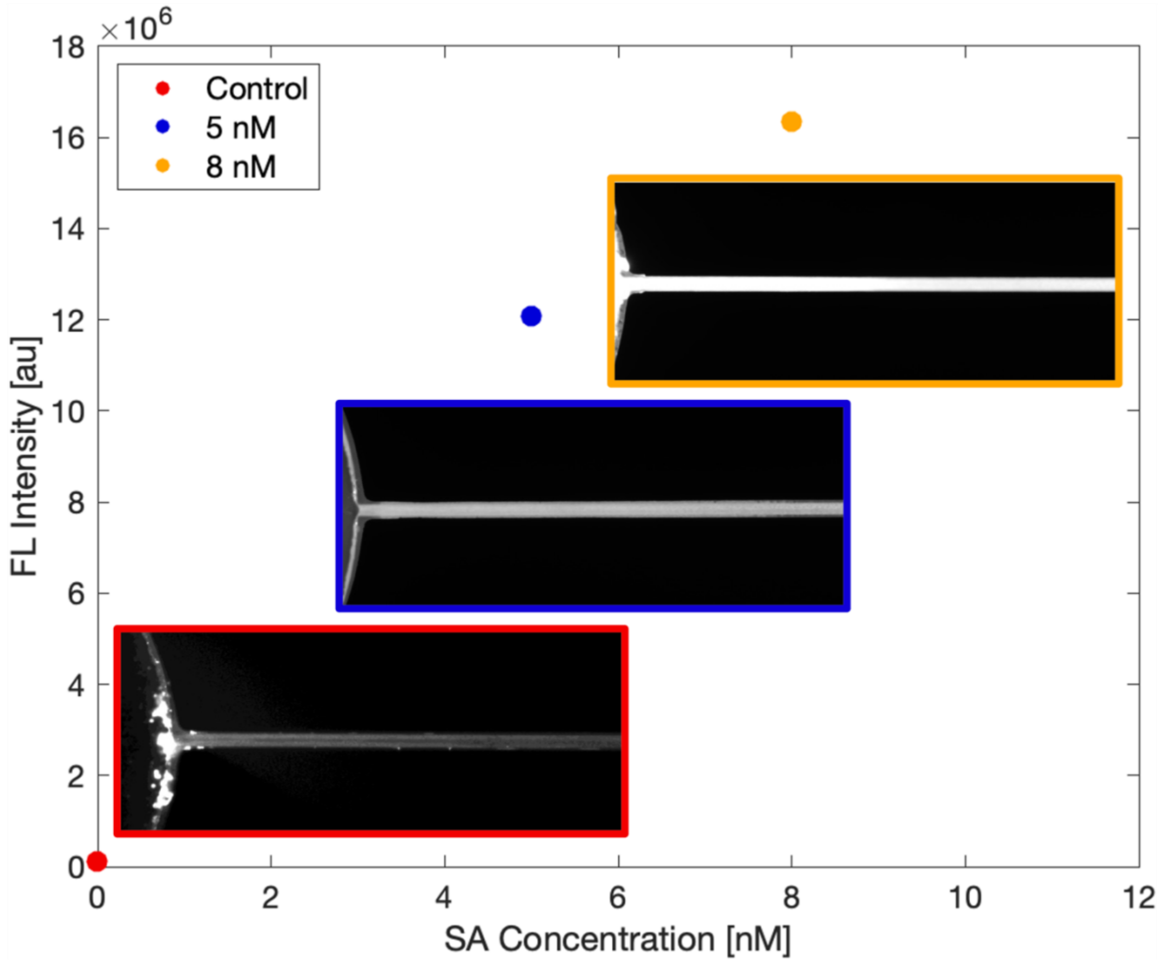


Figure *S9 – Fluorescent intensity of SA on the enclosed microchip*

### S11. Non-uniform coverage of the FL-tagged SA on the enclosed microchip

Due to suboptimal cleaning of the enclosed microchip, the surface is not uniformly activated by the RCA1 cleaning. Therefore, affinity probes do not uniformly cover the microchannels. Figure S10 shows the non-uniform coverage of FL-tagged SA inside the microchannels and uniform coverage in case of the open-top microchip. The fluorescent intensity drops significantly the further it gets from the fluidic ports in the enclosed microchip where the cleaning solution could diffuse and activate the surface prior to the measurements. The bottom inset of the figure shows the uniform FL intensity from the immobilized SA on the open microchip.

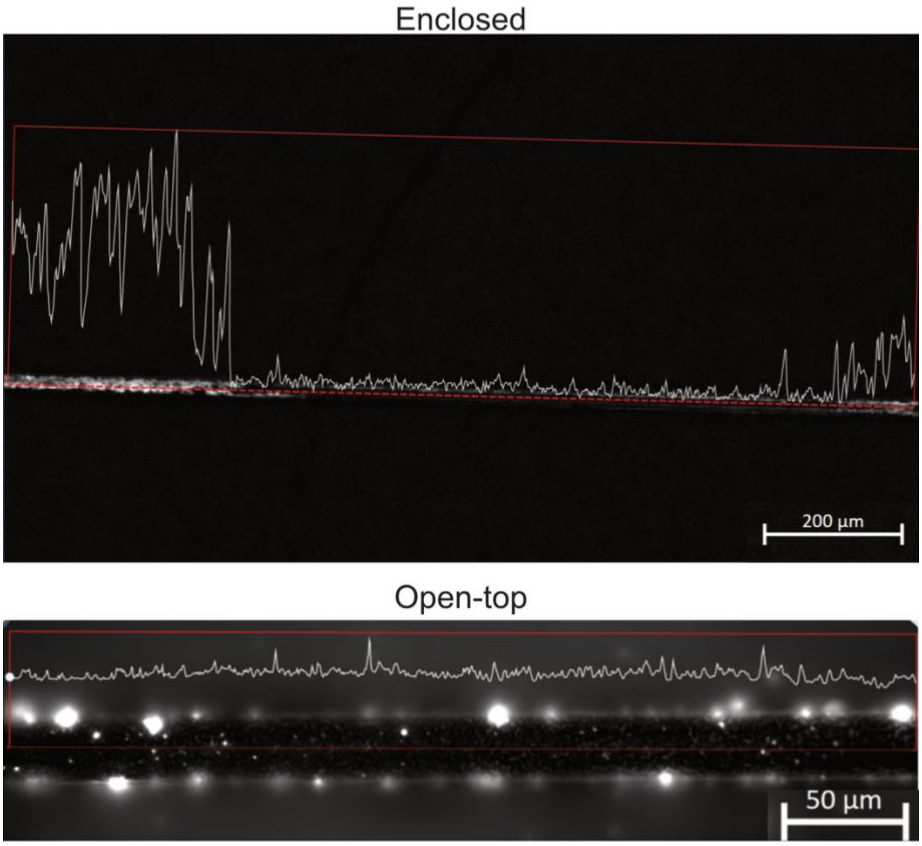


*Figure S10 –coverage of the microchannel by FL tagged SA in the enclosed microchip (top) and open-top (bottom)*

### S12. NTA size characterization of the sEVs

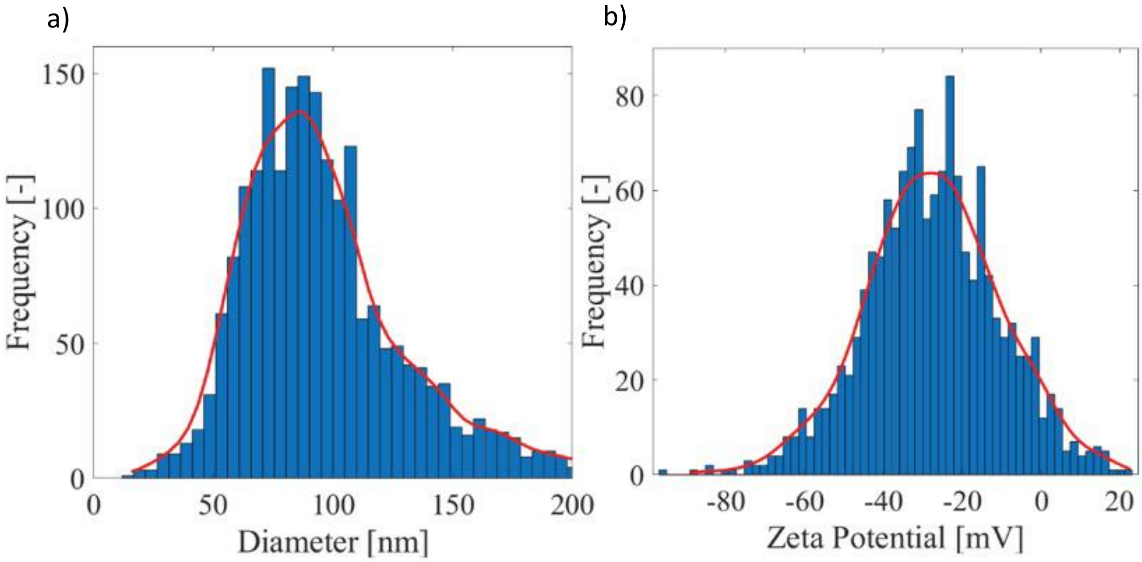


Figure *S11 – a) size distribution of the EVs, b) zeta potential distribution of the sEVs*

### S13. Control measurements on PPB and SPB devices

For the control measurements, an open-top microchip was functionalized up to the antibody immobilization for both PPB and SPB cases. Then 0.01 w% Pluronic-F108 was incubated on the surfaces for 30 minutes to passivate the uncovered surfaces. After washing the first baseline was measured. Then the highest concentration of sEVs (1x$10^8$ sEV/mL) was incubated on the microchips and the second baseline was measured.  As Figure S12 shows, the control signal on both surfaces is close to the MDS (5 pA). The control signal was subtracted from the sEV sensing data presented in the manuscript.

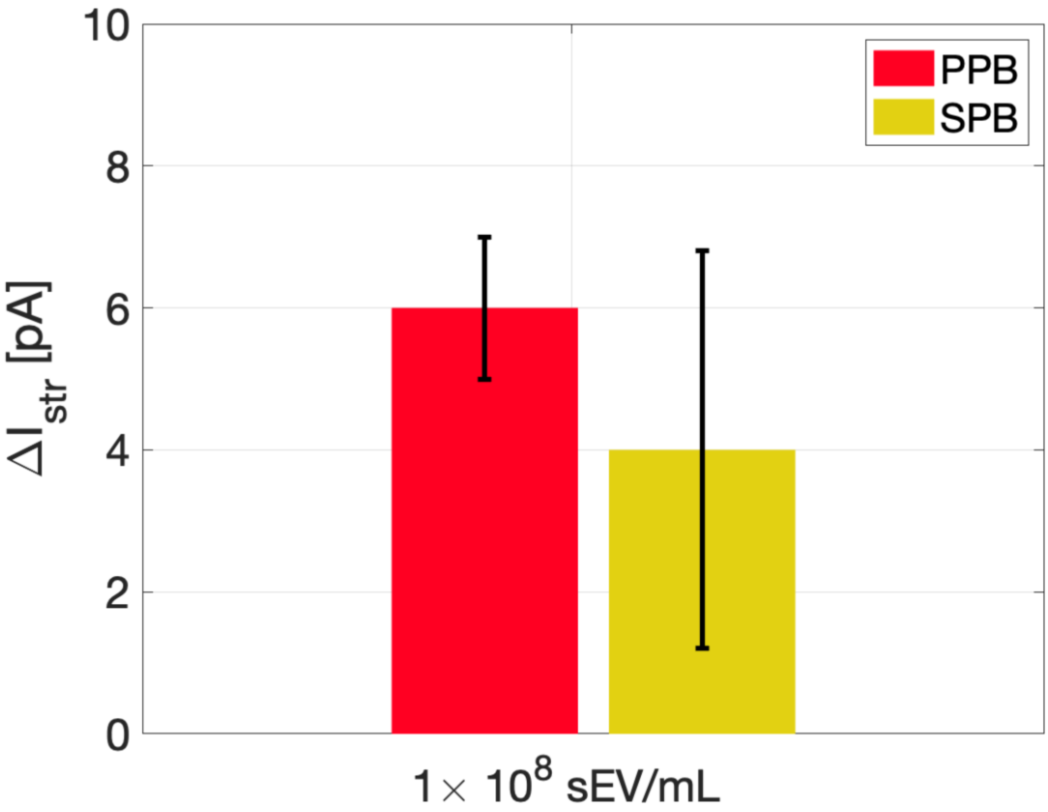


*Figure S12 – Control measurements for SPB and PPB surfaces at the highest concentration of sEVs*

**S14. Slide angle measurement**

To estimate the friction force between the liquid and PPB/SPB surfaces, two flat silica coverslips were functionalized by PPB and SPB following the protocol explained in the manuscript. Thereafter a 4 uL droplet of 0.1x PBS was placed on the center of the surfaces. The slope of the surfaces was gradually increased until the droplets started sliding along the surfaces. An image was snapped at this movement and the friction force was estimated as shown in Figure S13.

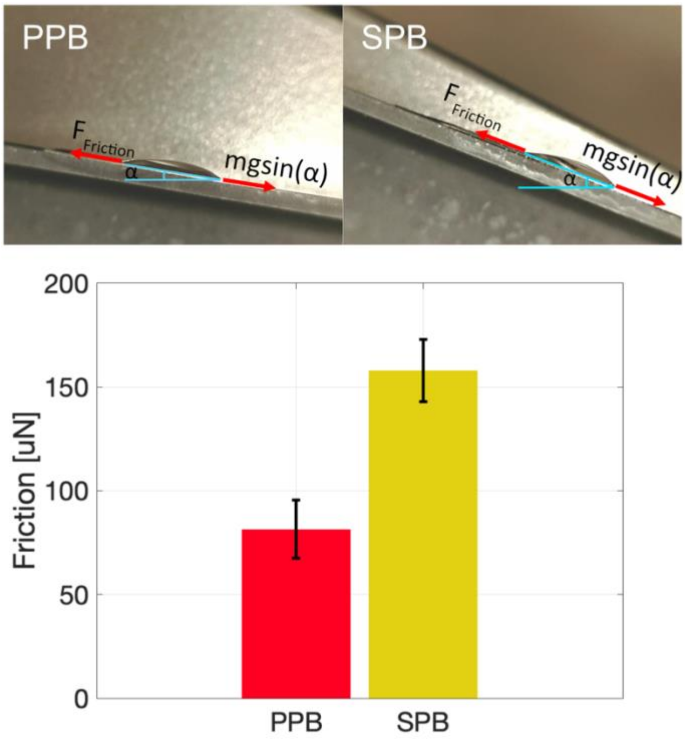


*Figure S13 – Slide angle measurement and friction force estimation between the liquid and the surface on both PPB and SPB surfaces*

**References:**


[1] C. Stiller *et al.*, “Detection of tumor-associated membrane receptors on extracellular vesicles from non-small cell lung cancer patients via immuno-PCR,” *Cancers (Basel).*, vol. 13, no. 4, p. 922, 2021.

[2] S. S. Sahu *et al.*, “Electrokinetic sandwich assay and DNA mediated charge amplification for enhanced sensitivity and specificity,” *Biosens. Bioelectron.*, vol. 176, p. 112917, 2021.

[3] S. S. Sahu *et al.*, “Multi-marker profiling of extracellular vesicles using streaming current and sequential electrostatic labeling,” *Biosens. Bioelectron.*, vol. 227, p. 115142, 2023.

[4] Z. Adamczyk, M. Nattich, and M. Zaucha, “Electrokinetics of particle covered surfaces,” *Curr. Opin. Colloid Interface Sci.*, vol. 15, no. 3, pp. 175–183, 2010.
